# Carvedilol suppresses ryanodine receptor-dependent Ca^2+^ bursts in human neurons bearing *PSEN1* variants found in early onset Alzheimer’s disease

**DOI:** 10.1101/2023.09.15.558029

**Authors:** Atsushi Hori, Tomohiko Ai, Takashi Hato, Haruka Inaba, Kimie Tanaka, Shoichi Sato, Mizuho Okamoto, Yuna Horiuchi, Faith Jessica Paran, Yoko Tabe, Corina Rosales, Wado Akamatsu, Takashi Murayama, Nagomi Kurebayashi, Takashi Sakurai, Takashi Miida

**Author notes:** Corresponding Author: Tomohiko Ai, MD, PhD.

## Abstract

Seizures are increasingly being recognized as the hallmark of Alzheimer’s disease (AD). Neuronal hyperactivity can be a consequence of neuronal damage caused by abnormal amyloid β (Aß) depositions. However, it can also be a cell-autonomous phenomenon causing AD by Aß-independent mechanisms. Indeed, various studies using animal models showed that Ca^2+^ releases from the endoplasmic reticulum (ER) via type 1 inositol triphosphate receptors (InsP_3_R1s) and ryanodine receptors (RyRs). To investigate which is the main pathophysiological mechanism in human neurons, we measured Ca^2+^ signaling in neural cells derived from three early-onset AD patients harboring variants of Presenilin-1 (*PSEN1* p.A246E, p.L286V, and p.M146L). Of these, it has been reported that PSEN1 p.A246E and p.L286V did not produce a significant amount of abnormal Aß. We found that all *PSEN1*-mutant neurons, but not wild-type, caused abnormal Ca^2+^-bursts in a manner dependent on the calcium channel, Ryanodine Receptor 2 (RyR2). Indeed, carvedilol, anRyR2 inhibitor, and VK-II-86, an analog of carvedilol without the β-blocking effects, sufficiently eliminated the abnormal Ca^2+^ bursts. In contrast, Dantrolene, a RyR1 inhibitor, and Xestospongin c, an IP_3_R inhibitor, did not attenuate the Ca^2+^-bursts. The RNA-Seq data revealed that ER-stress responsive genes were increased, and mitochondrial Ca^2+^-transporter genes were decreased in PSEN1_A246E_ cells compared to the WT neurons. Thus, we propose that aberrant Ca^2+^ signaling is a key link between human pathogenic *PSEN1* variants and cell-intrinsic hyperactivity prior to deposition of abnormal Aß, offering prospects for the development of targeted prevention strategies for at-risk individuals.

**One Sentence Summary:** Aberrant Ca^2+^-signaling causes *PSEN1*-related early onset Alzheimer’s disease.

## INTRODUCTION

Epileptic activities in Alzheimer’s (AD) patients have been gaining attentions due to the potential association with their abnormal cognitive behaviors[1, 2]. Specifically, seizures tend to appear more frequently in patients with early-stage AD than in elderly patients[1]. Indeed, in 2017, Lam et al. recorded subclinical seizure-like local neural hyper-excitations using intracranial catheters placed adjacent to the hippocampus in two human AD patients. The study showed that elimination of the subclinical seizures with levetiracetam, an anticonvulsant, improved the patient’s abnormal cognitive behavior although the underlying mechanisms of the seizure-like signals remain unknown[3]. Therefore, several different types of anticonvulsants have been used to treat AD. However, there is no clear consensus on which medications are suitable for which patients’ symptoms, such as epileptic amnesia and cognitive decline since the pathophysiological mechanisms of seizures in AD remains unelucidated [2].

To date, several hundreds of variants in the presenilin 1 gene (*PSEN1*) have been linked to early-onset familial Alzheimer’s disease (AD)[4, 5]. PSEN1 located in the membrane of the endoplasmic reticulum (ER) regulates intracellular Ca^2+^ homeostasis [5]. It has been reported that *PSEN1* variants found in AD produce abnormal amyloid β (Aß) and Tau proteins that damage neurons [6, 7]. However, recent studies revealed that not all pathogenic *PSEN1* variants produce abnormal Aß proteins [8], illustrating the importance of amyloid-independent mechanisms in the development of AD [9].

Also, intracellular Ca^2+^-dysregulation has been observed in various AD animal models. In a classic murine AD model bearing a *PSEN1* p.M146V variant, Glutamate provoked abnormal persistent Ca^2+^ waves probably via the NMDA receptors, which may have an association with neurodegeneration [10]. Shilling et al. proposed that the type 1 inositol triphosphate receptor (InsP_3_R1) is a main mediator to release Ca^2+^ from the ER in the *PSEN1* p.M146V knock-in mouse model [11]. Yao et al. reported that reduction of Ca^2+^-release via a mutated ryanodine receptor 2 (RyR2 p.E4872Q) from the ER prevented abnormal behaviors, such as hyperactivity, hyperexcitability, and memory loss in a severe early-onset AD mouse model (5xFAD), suggesting the role of RyR2 in Ca^2+^ dysregulation [12]. Based on these results, Schrank et al. generated a human neuronal model using iPS cells (iPSCs) obtained from an AD patient. They observed that production of abnormal Aß_42_ and abnormal Ca^2+^ were more prominent in the AD neurons than control cells, and dantrolene, an inhibitor of RyR,1 eliminated both Aß_42_ and Ca^2+^ surges [13]. However, these data still support the classical amyloid-theory, which may not be the main pathological mechanism.

Therefore, we hypothesize that dysregulated Ca^2+^ homeostasis and resultant neuronal hyperexcitability, independent from the abnormal Aß-theory, could be a discernable feature of pathogenic *PSEN1* mutations. Accordingly, we analyzed intracellular Ca^2+^ activities in iPS cell-derived neurons from multiple early onset AD patients with pathogenic *PSEN1* variants that do not produce abnormal amyloid [8]. We found that all neuronal cells bearing *PSEN1* variants exhibited abnormal Ca^2+^-bursts, but not WT neurons. Also, the RNA-Seq data showed that in the PSEN1_A246E_ cells, ER stress responsive genes such as *EIF2S1* and *ATF4* were more expressed than the WT neurons. These data implicate a potential higher risk of subclinical seizures in this population.

## MATERIALS AND METHODS

### Cell Culture and Immunostaining Reagents

All cells used in this study were commercially available cell lines purchased from Axol Bioscience (Cambridge, United Kingdom). This study protocol has been approved by the Institutional Review Board for Medical Research at the Juntendo University School of Medicine (IRB#E23-0210). Please contact them by the following email if necessary. Obtaining informed consents from the patients was impossible since all cells were purchased from Axol Bioscience. Thus, the informed consents were waived by the IRB committee.

The reagents that were used in this study are listed as follows: ibidi GmbH, Gräfelfing, Germany: Polymer coverslip-bottom dishes (ib81156); chamber slide (ib80826); mounting medium with 4′,6-diamidino-2-phenylindole (DAPI) (ib50011). Axol Biosciences, Cambridge, UK: SureBond-XF (ax0053); Neural Maintenance Medium Supplement (ax0031); Neural Maintenance Basal Medium (ax0031); human FGF2 (ax0047); human EGF (ax0048); NeurOne Supplement A (ax0674a); NeurOne Supplement B (ax0674b). Merck, Darmstadt, Germany: polyornithine aqueous solution (P36555); bovine serum albumin (A2153); anti-β-tubulin Ⅲ antibody (T8660); anti-microtubule-associated protein2 antibody (AB5622); 2-Mercaptoethanol (M3148); ascorbic acid (A5960); N6,2′-O-dibutyryladenosine 3′,5′-cyclic monophosphate sodium salt (D0260). Thermo Fisher Scientific, Waltham, USA: B-27 supplement (17504044); GlutaMAX supplement (35050061); 488-labeled secondary antibodies (S11223), 594-labeled secondary antibodies (S11227); Neurobasal-A Medium, minus phenol red (12349015). Biolegend, San Diego, USA: human Brain-derived neurotrophic factor (788904). Alomone labs, Israel: anti-ryanodine receptor 2 antibody (ARR-002). Abcam, UK: anti-Presenilin 1 antibody (ab15458). Wako Pure Chemical Industries, Osaka, Japan: paraformaldehyde (160-16061); Polyoxyethylene (10) Octylphenyl Ether (160-24751). NACALAI TESQUE, Kyoto, Japan: penicillin and streptomycin (26253-84), VectorLaboratories, Burlingame, USA: normal goat serum (S-1000).

### Preparation of iPSC-derived neural stem cells (NSCs)

Four iPSC-derived NSCs were purchased from Axol Bioscience (Cambridge, UK). Information about the donors was described in the texts and is also available online (https://www.axolbio.com/). Briefly, PSEN1_A246E_ cell line was established from a 31-year-old female bearing *PSEN1* p.A246E with early onset AD (onset age at 45 years-old; AX0114) [14]. PSEN1_L286V_ cell line was established from a 38-year-old male bearing *PSEN1* p.L286V with early onset AD (AX0112). PSEN1_M146L_ cell line was established from a 53-year-old male bearing *PSEN1* p.M146L with AD (AX0113). As a control (WT), a cell line established from a 64-year-old healthy female subject (AX0019) was used. All cells were differentiated into mature neurons for 27 ± 2 days.

Axol Bioscience performed karyotyping of these cells before and after differentiation, and no differences were found. All reagents used for culturing were purchased from Axol Bioscience (Cambridge, UK). The expansion of neural stem cells and neural cell differentiation was performed according to the protocol published by Axol Bioscience (Human iPSC-Derived Neural Stem Cells System A, B, C and D Protocol version 5.0: https://www.cosmobio.co.jp/product/uploads/document/AXO_Protocol_Human_iPSC_Derived_Neural_Stem_Cells.pdf and Enriched Cerebral Cortical Neurons Derivation from Axol iPSC Neural Stem Cells User Guide V1.4: https://axolbio.com/publications/enriched-cerebral-cortical-neuron-protocol/). Briefly, a 6-well plate was coated with SureBond-XF for at least 4 hours for expansion cultures. Human iPSC-derived NSCs were seeded and expanded in the 6-well plate. The medium was replaced with Neural Maintenance Medium (containing the Neural Maintenance Medium Supplement) adding human FGF2 (a final concentration of 20 ng/mL) and human EGF (a final concentration of 20 ng/mL). Polymer coverslip-bottom dishes were coated with polyornithine aqueous solution (0.05 mg/mL) and SureBond-XF for neuronal differentiation cultures. Human iPSC-derived NSCs were seeded in the polymer coverslip-bottom dishes. From day 1 to day 6 of the culture, the medium was replaced with Neural Maintenance Medium containing NeurOne Supplement A every other day. In addition, Neural Maturation Basal Medium was prepared by adding 1 mL of B-27 supplement, 0.5 mL of GlutaMAX supplement, 25 μL of 2-Mercaptoethanol, and 0.5 mL penicillin and streptomycin (a final concentration of 1%) to 50 mL of Neurobasal-A Medium, minus phenol red. From day 7 to day 14 of the culture, the medium was replaced with Neural Maturation Basal Medium containing NeurOne Supplement B, human Brain-derived neurotrophic factor (BDNF) (a final concentration of 0.02 μg/mL), N6,2′-O-dibutyryladenosine 3′,5′-cyclic monophosphate sodium salt (a final concentration of 0.5 mmol/L), and ascorbic acid (a final concentration of 0.2 mmol/L) every other day. On the 15th day, the whole volume of the medium was replaced with Neural Maturation Basal Medium supplemented with BDNF (a final concentration of 0.02 μg/mL), N6,2’-O-dibutyryladenosine 3′,5′-cyclic monophosphate sodium salt (a final concentration of 0.5 mmol/L), and ascorbic acid (a final concentration of 0.2 mmol/L). After that, half of the medium was replaced every other day.

### Ca^2+^ imaging

The intracellular Ca^2+^ levels were measured using Cal-520 (21130, AAT Bioquest, Inc., Sunnyvale, USA) as previously described [15]. Briefly, iPSC-derived neurons on polymer coverslip-bottom dishes as mentioned in section “Preparation of iPSC-derived neural stem cells (NSCs)” were incubated with the Cal-520/AM (4 µM) at 37℃ with 5% CO_2_ for 30 min in Neural Maturation Medium. The polymer coverslip-bottom dishes were washed with a Tyrode buffer (in mM:140 NaCl, 5.4 KCl, 5 HEPES, 1.2 MgCl_2_, 1.8 CaCl_2_, 10 glucose, pH 7.4). Data acquisition and analysis were performed using AquaCosmos 2.0 (Hamamatsu Photonics, Hamamatsu, Japan). Ethylene glycol-bis(*ß*-aminoethyl ether)-N,N,N’,N’-tetra acetic acid (EGTA) (15214-34) was purchased from NACALAI TESQUE (Kyoto, Japan). Cyclopiazonic acid (030-17171), dantrolene (359-44501) and xestospongin c (244-00721) were purchased from FUJIFILM Wako Chemicals (Osaka, Japan). Carvedilol (C184625) was purchased from Toronto Research Chemicals (Toronto, Canada). VK-Ⅱ-86 (AOB4007) was purchased from AOBIOUS INC (Gloucester, USA).

### Immunocytochemical staining

iPSC-derived neurons cultured in a chamber slide were fixed with 4 % paraformaldehyde at room temperature for 10 min. After permeabilization with 0.1% Triton X-100, sections were placed in a blocking buffer (5% goat serum and 2% bovine serum albumin in PBS) for 30 min, then labeled with primary antibodies with anti-β-tubulin Ⅲ antibody (1:500), anti-microtubule-associated protein2 antibody (1:500), anti-ryanodine receptor 2 antibody (1:300) and anti-Presenilin 1 antibody (1:50) overnight in blocking buffer in a humidified chamber at 4°C. For secondary reactions, Alexa 488-, 594-labeled secondary antibodies were used. Mounting medium with 4′,6-diamidino-2-phenylindole (DAPI) was used for nuclear staining. Fluorescence images were recorded with a TCS SP5 confocal microscope (Leica Camera Inc., Allendale, USA).

### RNA-Seq

RNA-Seq was performed as described anywhere [16]. Briefly, differentiated neurons at 28 days were rinsed with cold PBS and scraped using the lysis buffer (LRT) containing β-mercaptoethanol (#135-07522, Wako Chemicals, Osaka, Japan) on ice. RNA extraction was done using RNeasy Plus Mini Kit (Qiagen, 74134). Genomic DNAs were removed using gDNA Eliminator spin columns (Catalog #74134, Qiagen, Hilden, Germany). cDNAs were prepared with Clontech Smart-seq v4 ultra-low kit (Cat# 634888, Takara Bio, Shiga, Japan). PolyA tail enrichment was done using NEBNext® Poly(A) mRNA Magnetic Isolation Module (Cat No. E7490). Library preparation and sequencing were performed at Rhelixa using NEBNext® Ultra™II Directional RNA Library Prep Kit (Cat No. E7760). Approximately 15-20 M reads per library were generated using the Illumina NovaSeq 6000 sequencer. The sequenced data were mapped to GRCh38 genome (gencode40) using STAR. Uniquely mapped sequencing reads were assigned to GRCh38 gencode40 genes using featureCounts, and counts data were analyzed using edgeR.

### Statistics

Ca^2+^ signal data were classified into five categories (no signals, narrow wave, wide wave, oscillation, and burst) and their composition ratios were shown using GraphPad Prism software (GraphPad Software, San Diego, USA). Multiple comparisons of population proportions for each category were performed by Tukey’s test. Pixel intensity comparisons of PSEN1 and RYR2 in fluorescent immunostained images were performed by Mann-Whitney U test. *p*<0.05 was considered as statistically significant. All statistical analyses were performed using EXCEL Toukei Ver.7.0 (ESUMI Co., Ltd, Tokyo, Japan) and EZR (Saitama Medical Center, Jichi Medical University, Saitama, Japan) [17].

## RESULTS

### Differentiation of the iPSCs into neural cells

iPSC-derived neural stem cells (NSCs) from a healthy individual and early-onset AD patients bearing *PSEN1* variants were differentiated into neural cells as described in Methods. Differentiation of neural cells was confirmed by the expression of neuronal markers βIII-tubulin and microtubule associated protein 2 (MAP2) (**Fig 1**).

**Fig 1.**
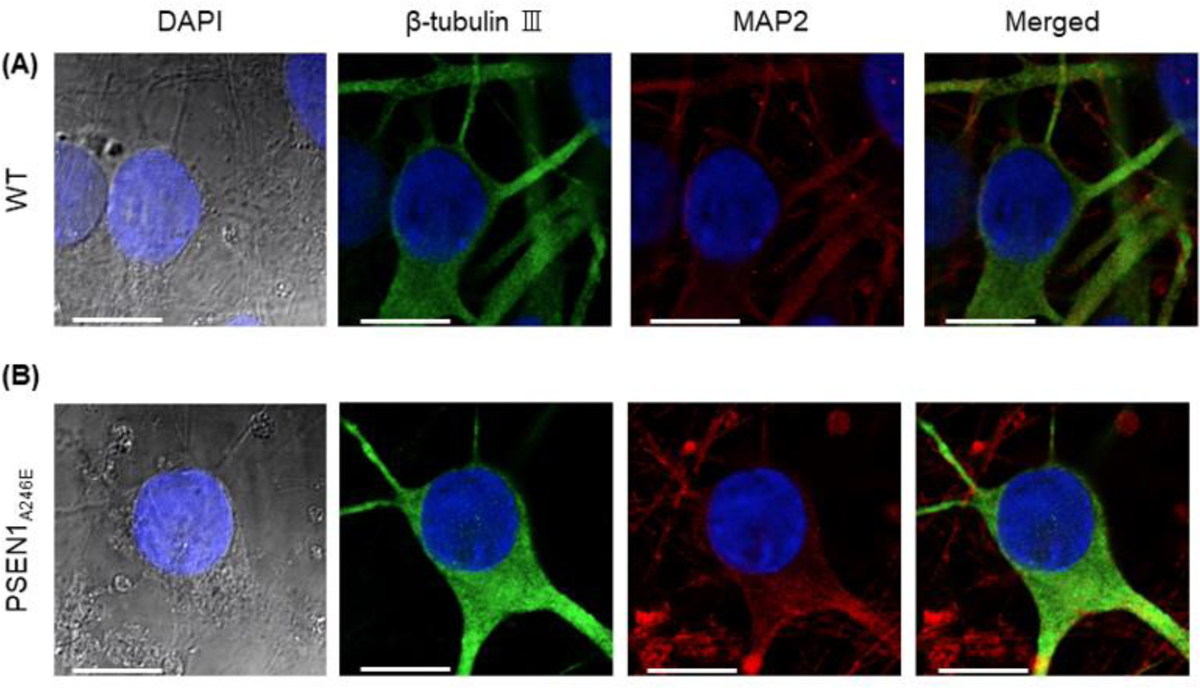
Immunocytochemical staining of WT and PSEN1 p.A246E neurons. Images obtained from WT neurons (A) and PSEN1 p.A246E neurons (B). The far-left panels are negative controls without using primary antibodies. Blue, DAPI; Red, β-tubulin Ⅲ; Green, MAP2; Scale bar, 10 µm.

### The iPSC-derived neurons bearing *PSEN1* variants exhibited abnormal Ca^2+^-waves

We found that all three types of neurons bearing *PSEN1* variants (p.A246E, p.M146L, and p.L286V) showed abnormal Ca^2+^-waves, most notably Ca^2+^-oscillations and bursts. In contrast, WT neurons showed occasional spontaneous Ca^2+^-waves (**Fig 2A-G**). **Fig 2H** summarizes the frequency of different Ca^2+^-wave patterns observed in each cell type. The abnormal Ca^2+^-oscillations were more frequently observed in the neurons bearing *PSEN1* variants (p.A246E, p.M146L, and p.L286V) compared to the WT neurons. Furthermore, Ca^2+^-bursts were significantly higher in neurons bearing *PSEN1* variants (p.A246E and p.M146L) compared to the WT neurons, suggesting that these two variants may cause more severe phenotypes than p.L286V since Ca^2+^-bursts are more disordered than Ca^2+^-oscillations.

**Fig 2.**
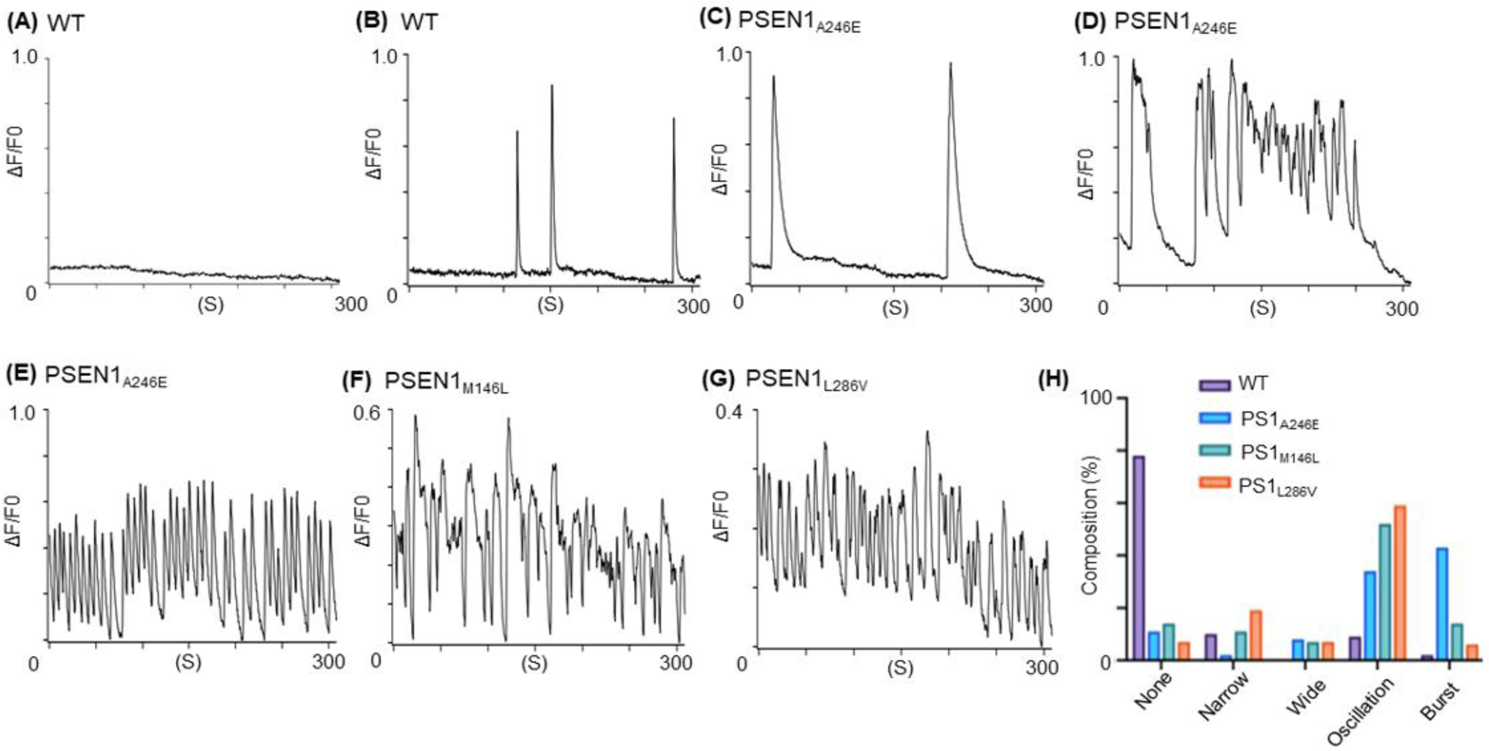
Representative traces of Ca^2+^ transients and their frequencies observed in various neuronal cells. (**A**) No Ca^2+^ transients observed in a WT neuron. (**B**) Narrow Ca^2+^ transients observed in a WT neuron. (**C**) Wide Ca^2+^ transients observed in a PSEN1_A246E_ neuron. (**D**) Ca^2+^-oscillations observed in a PSEN1_A246E_ neuron. (**E**) Ca^2+^-bursts observed in a PSEN1_A246E_ neuron. (**F**) Ca^2+^-bursts observed in a PSEN1_M146L_ neuron. (**G**) Ca^2+^-bursts observed in a PSEN1_L286V_ neuron. (**H**) The bar graphs show the frequencies of different Ca^2+^ transient in WT and *PSEN1*-mutated (p.A246E, p.M146L, p.L286V) neuronal cells. Numbers of cell counts for WT and *PSEN1*-mutated neurons were as follows: WT, 592; PSEN1_A246E_, 353; PSEN1_M146L_, 155; and PSEN1_L286V_, 363. Ca^2+^ transients were recorded at a sampling rate of 370 ms.

### Involvement of endoplasmic reticulum (ER) in the abnormal Ca^2+^-bursts

In human neurons, intracellular Ca^2+^ homeostasis is regulated by Ca^2+^ influx via Ca^2+^ channels located in the plasma membranes and Ca^2+^-induced Ca^2+^ release (CICR) from the ER via RYR and inositol 1,4,5-trisphosphate (IP_3_) receptors [18]. In addition, it has been proposed that PSEN1 variants may form pores in the ER membrane [19], leading to Ca^2+^ leaks into the cytoplasm through these pores [5]. To examine how the variant PSEN1 p.A246E causes Ca^2+^ leaks, the cells were perfused with Ca^2+^-free extracellular solutions containing 10 mM EGTA. Removal of extracellular Ca^2+^ eliminated the abnormal Ca^2+^ - bursts (**Fig 3A**). This indicates that abnormal Ca^2+^-waves associated with the PSEN1 p.A246E are CICR-dependent.

**Fig 3.**
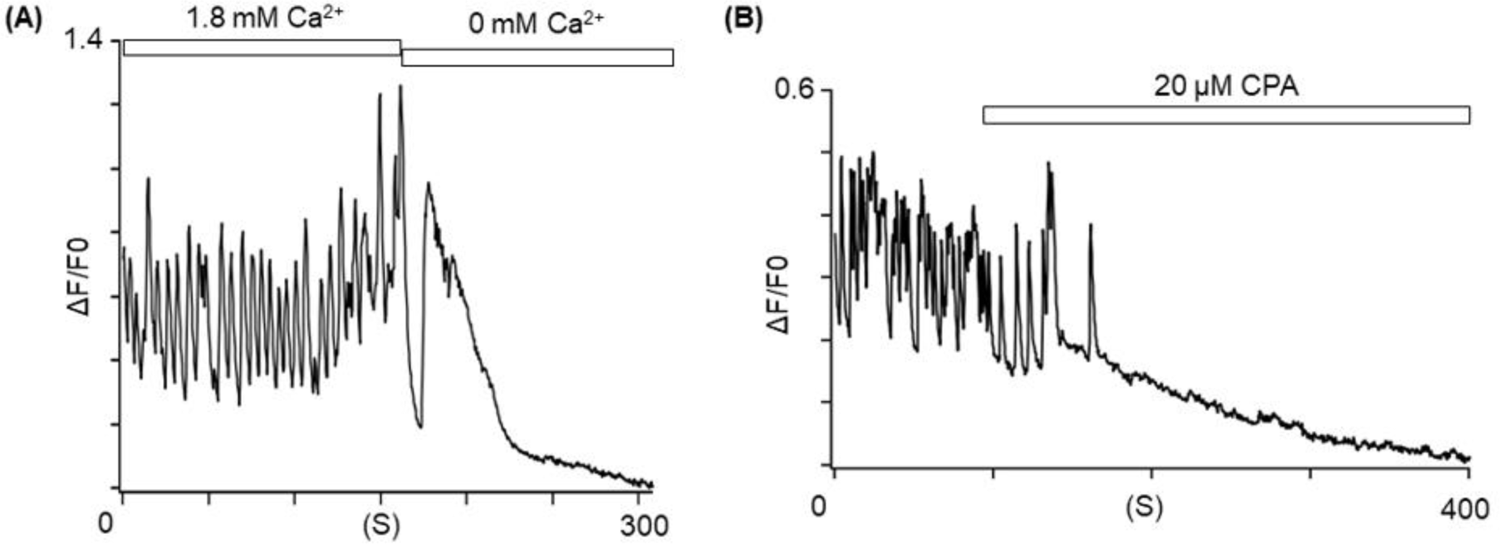
Involvement of endoplasmic reticulum in the underlying mechanisms of Ca^2+^ bursts in the PSEN1_A246E_ neurons. (**A**) Ca^2+^-free extracellular solutions containing EGTA eliminated the Ca^2+^-bursts in the PSEN1_A246E_ neuron. (**B**) Application of 20 μM CPA eliminated the Ca^2+^-bursts in the PSEN1_A246E_ neuron (N = 27).

To examine whether the ER is involved, cyclopiazonic acid (20 μM), an inhibitor of sarco/endoplasmic reticulum Ca^2+^-ATPase (SERCA), was applied to the Ca^2+^-waves [20], and it had suppressed the abnormal Ca^2+^-transients in the cells. Since cyclopiazonic acid depletes the Ca^2+^ storage in the ER, these results indicate that the ER is indeed the main resource of the abnormal Ca^2+^ bursts (**Fig 3B**).

### Ryanodine receptor (RyR) 2 facilitates the abnormal Ca^2+^-bursts

It has been reported that inositol triphosphate receptor (IP_3_R) is a mediator to release Ca^2+^ from the ER [11]. To examine the involvement of IP_3_R in the abnormal Ca^2+^-bursts [18], xestospongin c (3 μM), an IP_3_R antagonist, was applied to the Ca^2+^-bursts in PSEN1_A246E_ cells, which did not eliminate the abnormal Ca^2+^-bursts (**Fig 4A**).

**Fig 4.**
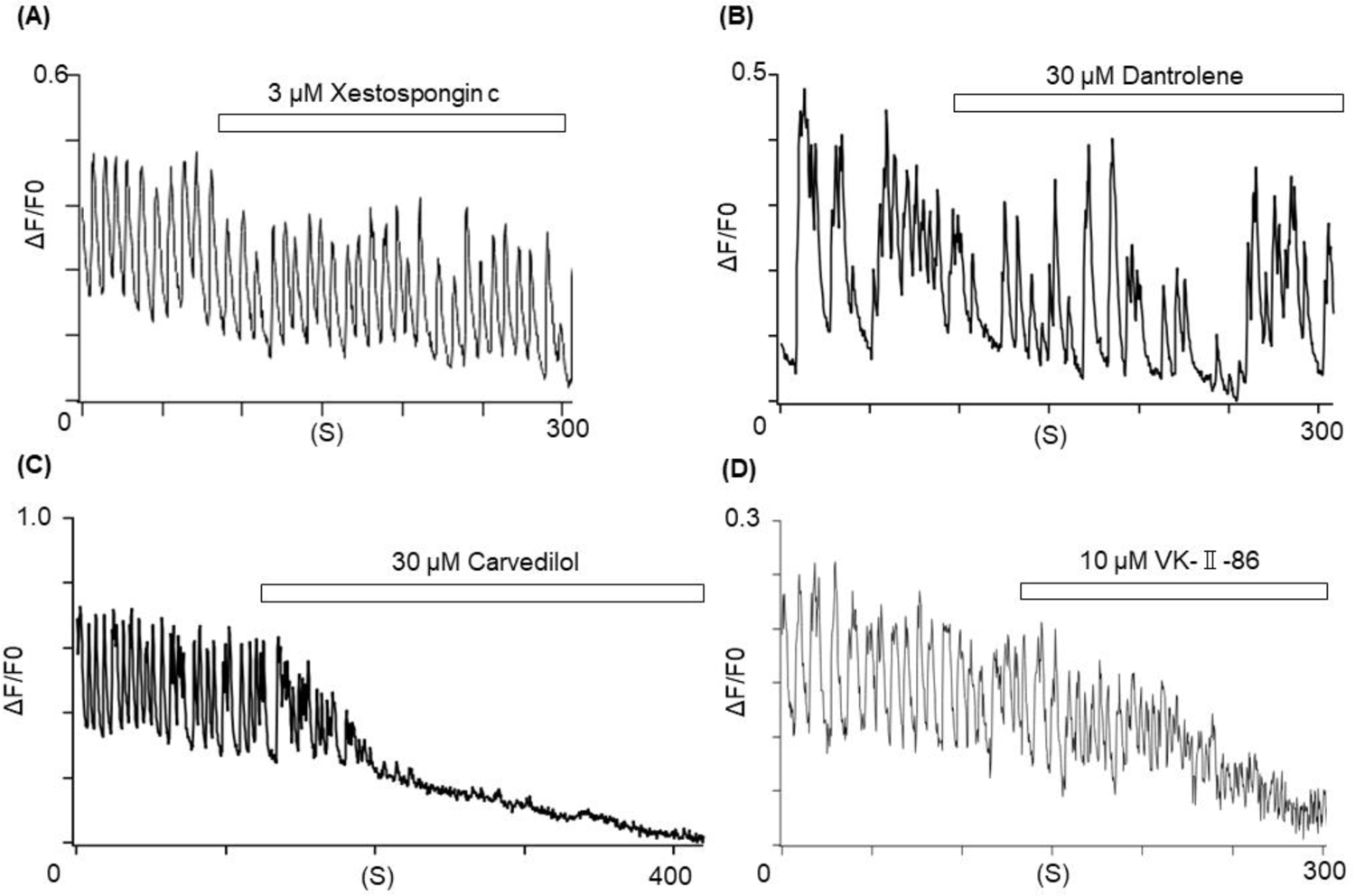
RYR2 facilitates the Ca^2+^ bursts from the ER in the PSEN1_A246E_ neurons. (**A**) Xestospongin c (3 μM) did not affect the Ca^2+^-bursts in the PSEN1_A246E_ neuron (N = 20). (**B**) Dantrolene (30 μM) did not affect the Ca^2+^-bursts in the PSEN1_A246E_ neuron (N = 21). (**C**) Carvedilol (30 μM) terminated the Ca^2+^-bursts in the PSEN1_A246E_ neuron (N = 98). (**D**) VK-Ⅱ-86 (10 μM) terminated the Ca^2+^-bursts in the PSEN1_A246E_ neuron (N = 18).

Human neurons express RyR1and RyR2 that pump out Ca^2+^ into the cytoplasm from the ER with a predominance of RyR2[21]. We examined effects of dantrolene, an inhibitor of RyR1 [22], and carvedilol, an inhibitor of RyR2 [23], on the abnormal Ca^2+^-bursts in the neurons bearing *PSEN1* variants. We found that carvedilol (30 μM), but not dantrolene (30 μM), eliminated the abnormal Ca^2+^-transients in the neuronal cells bearing *PSEN1* p.A246E (**Fig 4B, C**). The involvement of RyR2 was also confirmed with VK-II-86 (10 μM), a RyR2 inhibitor without the β-blocking effects (**Fig 4D**) [23]. The Ca^2+^-bursts were also eliminated by carvedilol in the PSEN1_M146L_ and PSEN1_L286V_ cells (**Fig S1**). These results indicate that RyR2 is the main facilitator of the abnormal Ca^2+^-bursts in the cells bearing *PSEN1* variants.

### The expression levels of RyR2 were not affected by *PSEN1* p.A246E

In a presenilin-knock out mouse, the protein level of RyR was reduced, which affected the intracellular Ca-homeostasis [24]. It has also been reported that PSEN physically interacts with RyR2 [25]. To examine the effects of the PSEN1 variant on RyR2 levels, we quantified the levels of PSEN1 and RyR2 using immunocytochemical staining. Our data revealed that the intracellular distributions of RyR2 and PSEN1 were similar both in the WT and PSEN1_A246E_ cells, and these two proteins seem to be partially colocalized (**Fig 5A**). The expression levels of PSEN1 and RyR2 measured by the fluorescence intensities were comparable between the WT and PSEN1_A246E_ cells (**Fig 5B**).

**Fig 5.**
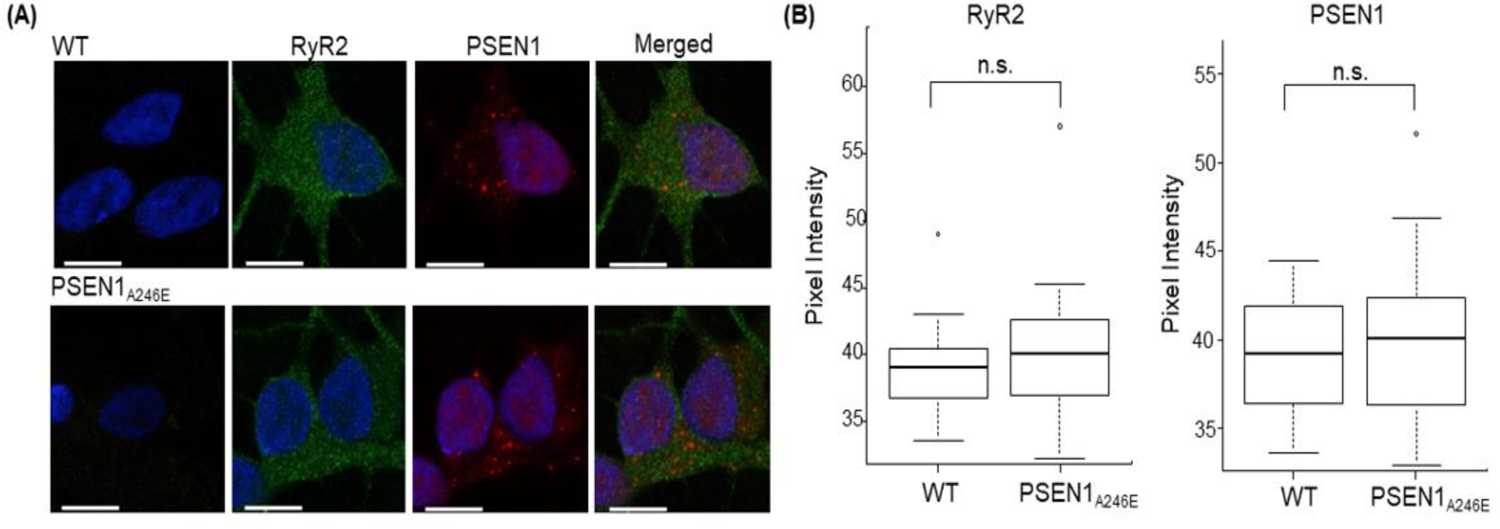
Immunocytochemical staining of RyR2 and PSEN1 in neural cells. **(A)** WT neurons (upper panels) and PSEN1 p.A246E (lower panels). Staining: blue, DAPI; green, RyR2; red, PSEN1. Scale bar, 10 µm. (**B**) Box plots of the intensity of PSEN1 in the WT (n = 33) and PSEN1_A246E_ (n =23) neurons (left panel). Box plots of the intensity of RYR2 in the WT (n = 33) and PSEN1_A246E_ (n = 23) neurons (right panel). All statistical analyses were performed by Mann-Whitney U test using EZR.

### RNA-Seq

In addition, the gene expressions were measured by RNA-Seq, and the results were compared between the WT and PSEN1_A246E_ neurons (n = 3 for each cell type). Genes with FDR *<* 0.05 were selected, recovering a list of the top 10000 differentially expressed genes (DEGs) with 1506 upregulated and 1521 downregulated in the PSEN1_A246E_ cells (**Table S1,** FDR < 0.05).

**Fig 6A** shows a smear plot of the alteration of the gene expression, annotated for genes associated with ER and mitochondrial-stress and glycolysis [26]. Among the top 10000 DEGs, the *MCUB* gene encoding for the mitochondrial calcium uniporter dominant negative subunit beta was upregulated, while the gene *MTLN* encoding mitoregulin and the gene *BCL2A1* encoding BCL2 related protein A1 were downregulated in the PSEN1_A246E_ cells. Also, the *RYR2* and *RYR3* genes encoding RyRs were downregulated in the PSEN1_A246E_ cells, which may reflect negative feedback of Ca^2+^ overload.

**Fig 6.**
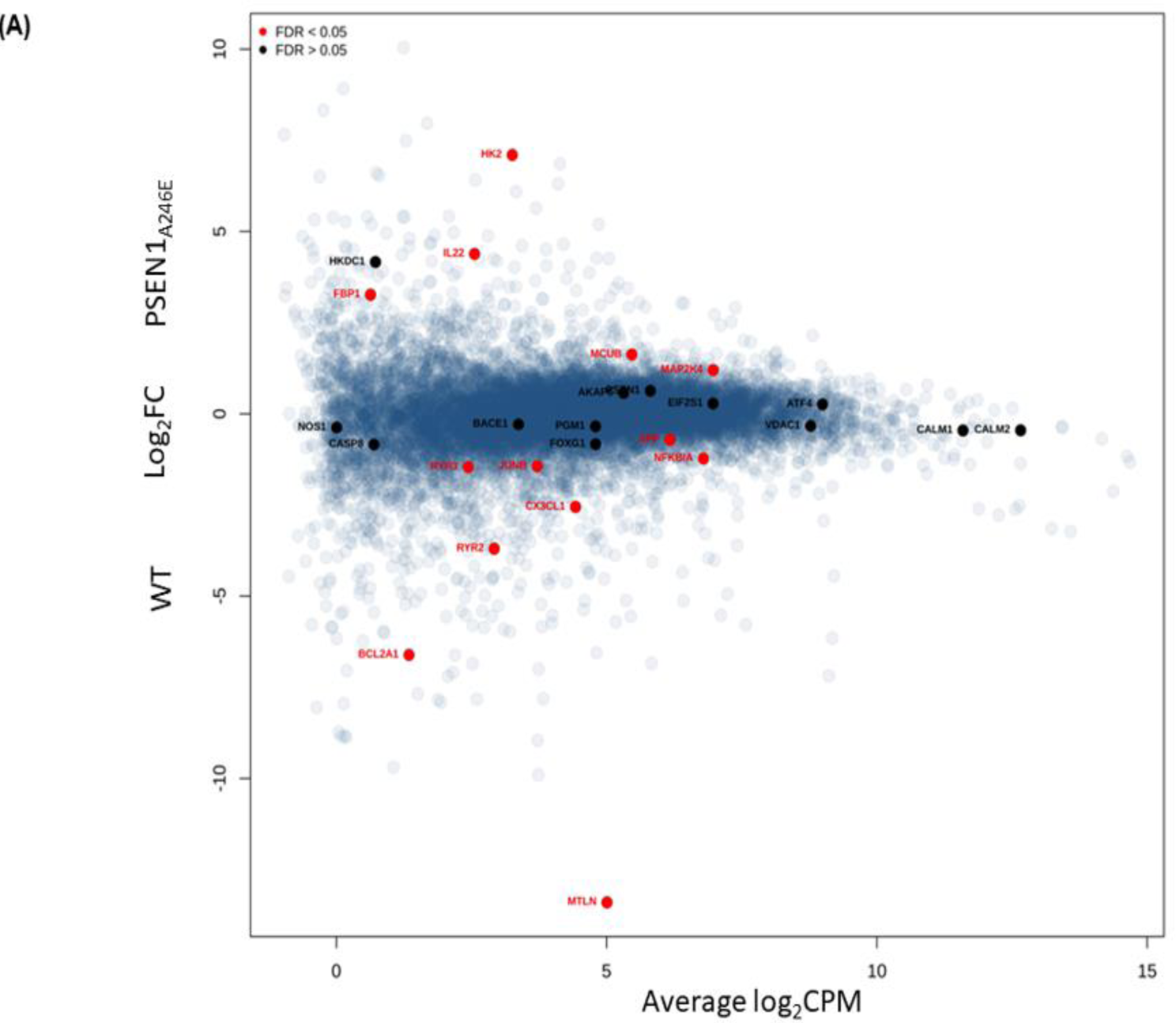
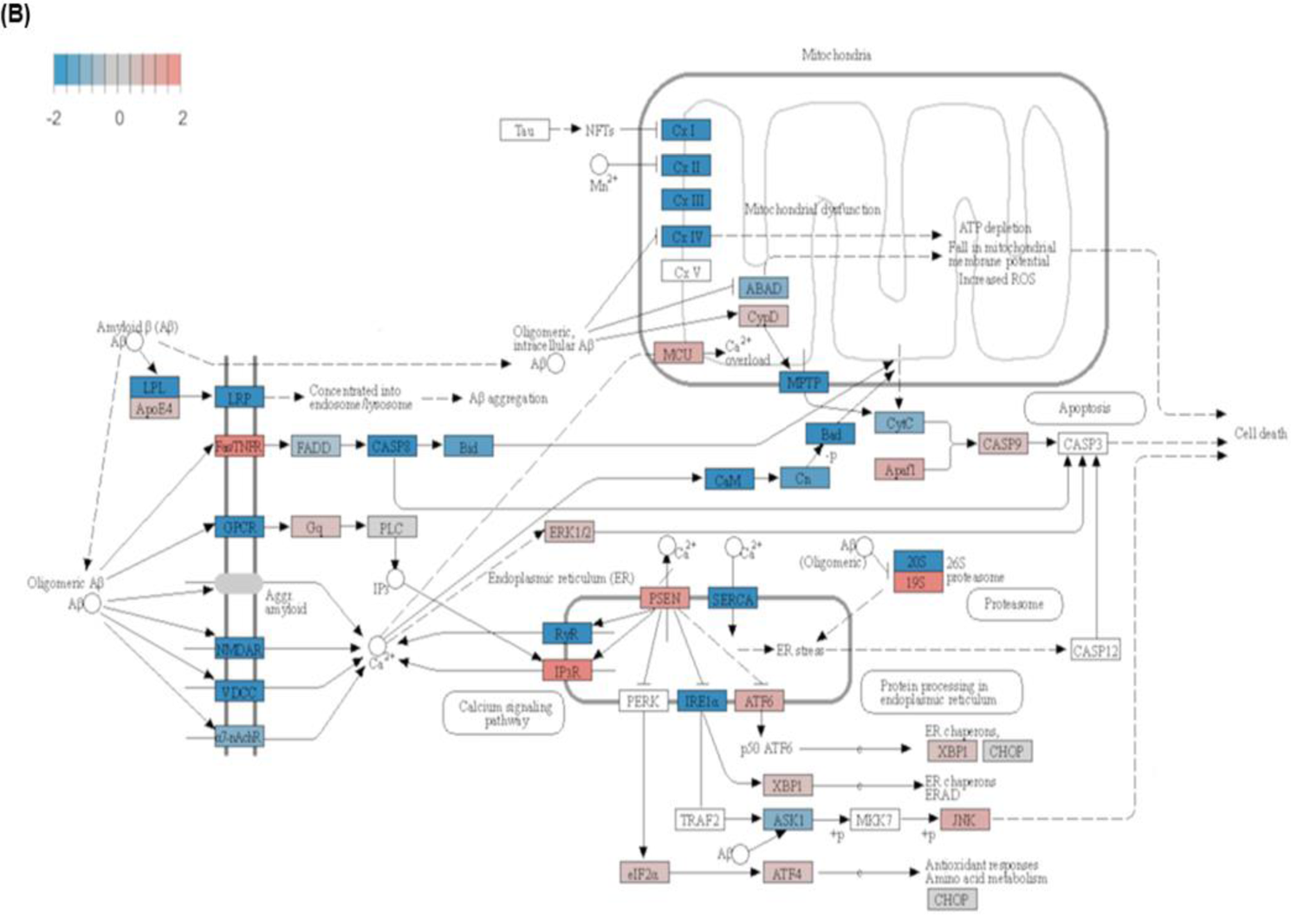
Comparisons of gene expressions between PSEN1_A246E_ and WT neurons evaluated by RNA-Seq. (A) A smear plot of detected gene expressions. Relative gene expression levels by comparisons of the PSEN1_A246E_ cells against the WT neurons are plotted. Several genes of interest were indicated by solid red symbols (FDR < 0.05). **(B)** Change of the gene expression in the ER and mitochondrial pathways associated with AD (partial KEGG Alzheimer’s Disease pathway map). Colors in the boxes indicate the genes tended to be downregulated (blue) or upregulated (red) in PSEN1_A246E_ cells, respectively.

The ER-processing pathway was one of the most significant enrichments. **Fig 6B** shows the pathway visualized using the Pathview package (KEGG hsa05010) [27], and the genes were labeled based on its fold change. The genes associated with ER-stress such as *EIF2S1* and *ATF4* tended to be upregulated in PSEN1_A246E_ neurons compared to the WT neurons (FDR > 0.05). These results suggest that the increase of Ca^2+^ release associated with *PSEN1* p.A246E may affect the ER and mitochondrial functions.

## DISCUSSION

Our data showed abnormal Ca^2+^-bursts recorded in human neuronal cells obtained from three-unrelated early onset AD patients bearing *PSEN1* variants (p.A246E, p.M146L, or p.L286V). We also demonstrated that the abnormal Ca^2+^-bursts were suppressed by carvedilol, a known RyR2 inhibitor, as well as VK-II-86, a RyR2 inhibitor without the β-blocking effects [23]. Importantly, these *PSEN1* variants do not produce significant amount of abnormal Aβ [8], which suggests that amyloid-independent pathophysiology plays a role in early-onset AD.

### Does neuronal hyperexcitation cause early-onset AD?

Although epileptic activity is often associated with AD, whether it is a cause, or a consequence of other factors, such as abnormal Aβ and Tau proteins, remains unknown [2]. However, seizures and myoclonus occur frequently in early-onset AD patients who have yet to develop severe dementia [1, 28]. This progression may be similar to pathological conditions of cardiomyopathy. In cardiomyopathy, it is often seen that abnormal electrical activities (i.e., arrhythmias) precede structural remodeling as how seizures progress to severe dementia [29, 30].

Just as Lam et al. proposed that silent seizure-like signals cause AD, our data strongly promotes a new underlying mechanism of *PSEN1*-associated early-onset AD. Since their patients did not bear *PSEN1* variants, the underlying mechanism of the neuronal hyperexcitation observed is unknown. However, *PSEN1* variants p.A246E and p.L286V tested in this study were found in AD patients who had seizures [31], which is notably consistent with Lam’s findings regardless of the causes of neuronal hyperexcitation. In addition, the *PSEN1* p.A246E and p.L286V as well as p.M146L variants did not produce significant amount of abnormal Aβ peptides [8], further supporting the amyloid-independent theory. Thus, it is reasonable to propose that neuronal hyperexcitation associated with Ca^2+^-dysregulation is one of the major amyloid-independent mechanisms of AD [3].

### The underlying mechanisms of Ca^2+^ dysregulation in *PSEN1*-related AD

There are numerous studies that Ca^2+^ overload plays an important role in damaging neurons though this topic is still controversial. For instance, Ca^2+^-leak from ER induces cell death by increasing the permeability of mitochondrial inner membranes [32]. Müller et al. proposed that constitutively activated transcription factor cAMP response element binding protein (CREB) associated with activation of IP_3_R through CaM kinase β and CaM kinase IV pathways cause cell death and sensitivity to Aβ toxicity [33]. In human-induced neurons from an AD patient bearing *PSEN1* p.A246E, increased Aβ_42_ and abnormal Ca^2+^ surge were observed and these were reversed by dantrolene [13], which is inconsistent with our data (**Fig 4B**). This can be due to differences in creating iPSCs and patient background.

Nonetheless, an important question moving forward is “How can we link local seizures due to abnormal Ca^2+^-bursts with cognitive behavioral disorders?” There are several suggestive reports. In an AD-mouse model (5xFAD) that bear multiple amyloid β precursor gene (*APP*) and *PSEN1* variants, reducing Ca^2+^ release from the ER via a mutated RyR2 prevented the hyperactivity of pyramidal neurons [12]. In another study, long-term treatment with carvedilol suppressed seizures in amyloid precursor protein transgenic mice (TgCRND8), a well-established AD model [34]. In contrast, Shilling et al. showed that, in a mouse model bearing *PSEN1* p.M146V, reduction of IP_3_R1 expression restored CREB-dependent gene expression rescuing the abnormal hippocampal long-term potentiation. Thereby, they propose that the IP_3_R1 is the therapeutic target in *PSEN1*-associated AD [11]. However, in our data, Xestospongin c, an inhibitor of IP_3_R, did not suppress the abnormal Ca^2+^-bursts (**Fig 4A**).

Our RNA-Seq data demonstrated that mitoregulin encoded by the *MTLN* that regulates mitochondrial membrane potential [35] and BCL2A1 encoded by the *BCL2A1* that is associated with apoptosis in brains [36] were significantly downregulated in the PSEN1_A246E_ neurons. In addition, ER stress pathway genes such as *EIF2S1* and *ATF4* [26], and mitochondrial stress pathway genes such as *MCU*, *CASP3,* and *CASP9* tended to be upregulated in the PSEN1_A246E_ compared to the WT neurons. In contrast, the *VDAC1* encoding the voltage-dependent anion channel 1 (VDAC1) in the outer mitochondrial membrane tended to be downregulated in the PSEN1_A246E_ [37]. Although these changes are not statistically significant (FDR > 0.05), it suggests that Ca^2+^-overload due to the abnormal Ca^2+^-bursts may damages cytosolic organelles, leading to neural malfunctions. Thus, currently, we are investigating the functional changes in the ER and mitochondria.

Though actual pathogenic mechanisms of AD are still controversial, it is reasonable to theorize that abnormal Ca^2+^-bursts through RyR2 can be one of the main pathophysiological mechanisms in AD associated with *PSEN1*.

### Limitations

There are several significant limitations: (1) we used human iPSC-induced neural cells whose anatomical and functional characteristics can be different from actual human neurons; (2) currently, it is difficult to generate homogeneous neural cell populations though we used a protocol to differentiate the cells into cerebral cortical neural cells; (3) although we examined the neural cells bearing three different *PSEN1* variants, we do not know if our hypothesis can be applied to AD caused by other factors such as *PSEN2* and *APP* variants; (4) we do not know the effects of these *PSEN1* variants on gating-kinetics of RyR2 due to lack of experiments in human brain cells. However, this is beyond the scope of this study; (5) our data still did not reveal how abnormal Ca^2+^-bursts cause early-onset AD (i.e., neural damages). Currently, we are studying the effects of Ca^2+^-bursts on ER and mitochondrial functions; (6) the RNA-Seq data showed that various pathways were affected by *PSEN1* p.A246E (listed in **Table S1**). Currently, we do not know the association of these changes and AD phenotypes. We will further investigate the roles of the affected various pathways in pathophysiological mechanisms of AD; and (7) a clinical trial failed to show the effects of carvedilol in AD (https://www.clinicaltrials.gov/study/NCT01354444?tab=history). Probably this was due to small population (n = 14), and we expect that carvedilol may not be able to reverse already damaged neurons. Interestingly, a recent large cohort study in Denmark indicated that the prevalence of AD was significantly lower in the patients who took carvedilol than the patients who took atenolol and bisoprolol [38]. However, it was unclear if the pathophysiology in these patients was uniform.

## Conclusions

Despite these limitations, we propose that abnormal Ca^2+^ signals play a role in the pathophysiological mechanisms in early-onset AD associated with *PSEN1*. Our data also elucidated that RyR2 can be a potential therapeutic target in prevention of AD. Further large-scale randomized clinical studies are warranted to validate our proposals.

## Acknowledgments

We thank the Laboratory of Molecular and Biochemical Research, Biomedical Research Core Facilities and the Laboratory of Morphology and Image Analysis, Research Support Center, Juntendo University Graduate School of Medicine, for technical assistance.

---

**Fig S1.**
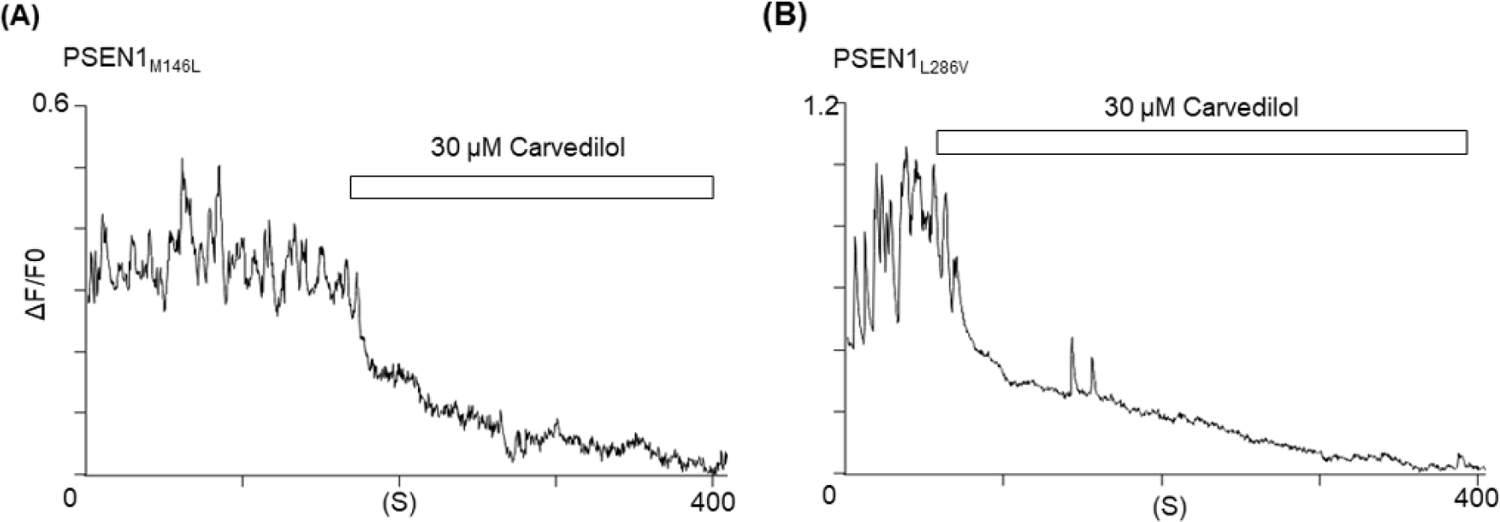
Effects of carvedilol on the Ca^2+^-bursts in the PSEN1_M146L_ and PSEN1_L286V_ neurons.

**Table S1.** A list of the top 1,000 genes that were affected in the PSEN1_A246E_ neurons compared to the WT neuron.

